# A back-translational study of descending interactions with central mechanisms of hyperalgesia induced by high frequency stimulation in rat and human

**DOI:** 10.1101/2022.11.25.517919

**Authors:** Ryan Patel, Joseph L Taylor, Anthony H Dickenson, Stephen B McMahon, Kirsty Bannister

## Abstract

In humans and animals, high frequency electrocutaneous stimulation (HFS) may produce an ‘early long-term potentiation-like’ sensitisation. Peripheral and central modulatory processes are proposed to play a role. To explore the impact of descending inhibitory pathway activation on the development of HFS-induced hyperalgesia, we concurrently applied HFS with i) a conditioned pain modulation (CPM) paradigm during psychophysical testing in humans, or ii) a diffuse noxious inhibitory controls (DNIC) paradigm during *in vivo* electrophysiological recording of spinal neurones in anaesthetised animals in parallel studies that utilised identical stimuli. HFS induced enhanced perceptual responses to pin-prick stimuli in cutaneous areas secondary to the area of stimulation in humans and heightened the excitability of spinal neurones in rats (which exhibited stimulus intensity dependent coded responses to pin-prick stimulation in a manner that tracked with human psychophysics), where we also observed indicators of increased central neuronal hyperexcitability. In humans, a HFS(+CPM) paradigm did not alter primary or secondary hyperalgesia, and the area and pain intensity of secondary hyperalgesia did not correlate with temporal summation of pain or CPM magnitude, while in rats application of a DNIC paradigm concurrent to HFS did not impact the development of neuronal hyperexcitability. Concordance between human and rat data supports their translational validity. Our finding that excitatory signalling exceeds inhibitory controls suggests that dampening facilitatory mechanisms may be a preferable strategy for certain chronic pain states. If facilitatory mechanisms dominate, our data could explain why enhancing activity in descending inhibitory controls is not sufficient to induce pain relief in vulnerable patients.

## 1. Introduction

Human surrogate models of pain have the potential to bridge the gap between pre-clinical and clinical research. Typically surrogate models are used to mimic the positive sensory phenomena associated with chronic pain states (Quesada et al., 2021). However, crucial to successful translation is the predictive value of animal models when assessing underlying mechanisms. Tetanic stimulation of C-fibres has long been known to elicit long term potentiation (LTP) of synaptic transmission in the rat dorsal horn (Liu and Sandkühler 1995; Randić et al., 1993), which is proposed as a prospective mechanism underlying activity dependent spinal plasticity in chronic pain states (Ikeda et al., 2006). ‘Early LTP-like’ phenomena have also been described in humans following high frequency stimulation (HFS), and the resultant primary and secondary hyperalgesia are hypothesised to reflect perceptual correlates of homosynaptic and heterosynaptic facilitation (Henrich et al., 2015; Klein et al., 2004).

From a functional perspective excitatory and inhibitory mechanisms likely exist in a counterbalanced manner to fine tune spinal transmission. Whilst spinal neuronal sensitisation can be heavily dependent on the nature of peripheral excitatory drive and subsequent synaptic plasticity, the overall resulting hyperexcitability will be determined by the relative strengths of these excitatory and inhibitory processes. Reversible spinal block of descending tracts can increase spinal neuronal excitability post-tetanic stimulation suggesting that descending inhibitions, in the context of inducing LTP, may act to restrict spatially and temporally the spread of sensitisation (Gjerstad et al., 2001). As excessive pain could arise from various combinations of increased excitatory and/or decreased inhibitory mechanisms at multiple levels throughout the sensory neuroaxis, this provides two contrasting strategies for pain management. It could then follow that central to the advancement of mechanistic-based pain medicine is not only the assessment of pain mechanisms but their interaction to develop rational pain treatment strategies.

In this study we examined whether applying conditioned pain modulation (CPM) concurrently to high frequency electrical stimulation affected the development of secondary hyperalgesia in healthy pain free subjects. Conditioned pain modulation is considered the psychophysical correlate of diffuse noxious inhibitory controls (DNIC) which is a unique form of pontospinal modulation, recruited by two distant noxious stimuli, mediated by the noradrenergic A5 nucleus via spinal α_2_-adrenoceptors (Bannister et al., 2015; Kucharczyk et al., 2022). By performing rat neurophysiological recordings alongside human psychophysics, we were able to investigate spinal and descending mechanisms during the application of identical test paradigms. Second order neurones in the deep dorsal horn encode multiple features of nociceptive processing including fine-tuned intensity coding (Maixner et al., 1986), spatial and temporal summation (Coghill et al., 1993; Mendell and Wall 1965), and unlike their superficial counterparts are modulated by DNIC (Le Bars et al., 1979a; b). Due to the convergence of peripheral and descending mechanisms upon these wide dynamic range neurones, they are aptly placed in the sensory pathway to provide a readout of neuronal substrates of peripheral and central sensitisation in a manner that relates to sensory testing measures to the same stimuli (O’Neill et al., 2015; Sikandar et al., 2013).

## 2. Methods

### 2.1 Subjects for the quantitative sensory testing study

Thirty seven healthy subjects were included in the study (eight male: 25.8 ± 5.03 years (range 20-41); twenty nine female: 25.4 ± 2.81 years (range 18-50)). Information about the study was disseminated both via the King’s College London fortnightly research volunteers email circular and within the department. Inclusion criteria specified that subjects should be between 18 and 70 years old, have a strong grasp of English, and be free of pain and medication (except contraception) on the day of testing. Exclusion criteria included acute or chronic pain conditions, dermatological issues at the site of testing, pregnancy, neurological disorders and musculoskeletal or inflammatory conditions. Subjects were requested not to consume alcohol in the 24 h period prior to testing, and to avoid excessively strenuous exercise of the legs. All subjects provided informed consent prior to testing which took place within the Wolfson Centre for Age Related Diseases. The study was approved by King’s College London Research Ethics Committee (Reference RESCM-21/22-22208) and performed according to the Helsinki Declaration (WMA 2013).

### 2.2 Protocol for quantitative sensory testing

Subjects attended two sessions which lasted approximately 2 hr. These sessions were set at least two days apart to avoid any confounding effects of long lasting sensitisation. During the HFS(control) session, subjects received HFS in isolation, whilst in the HFS(+CPM) session, subjects received HFS at the same time as a conditioning stimulus in the form of tonic pressure to the contralateral calf (Fig. 1A). Session order was randomised.

**Figure 1.**
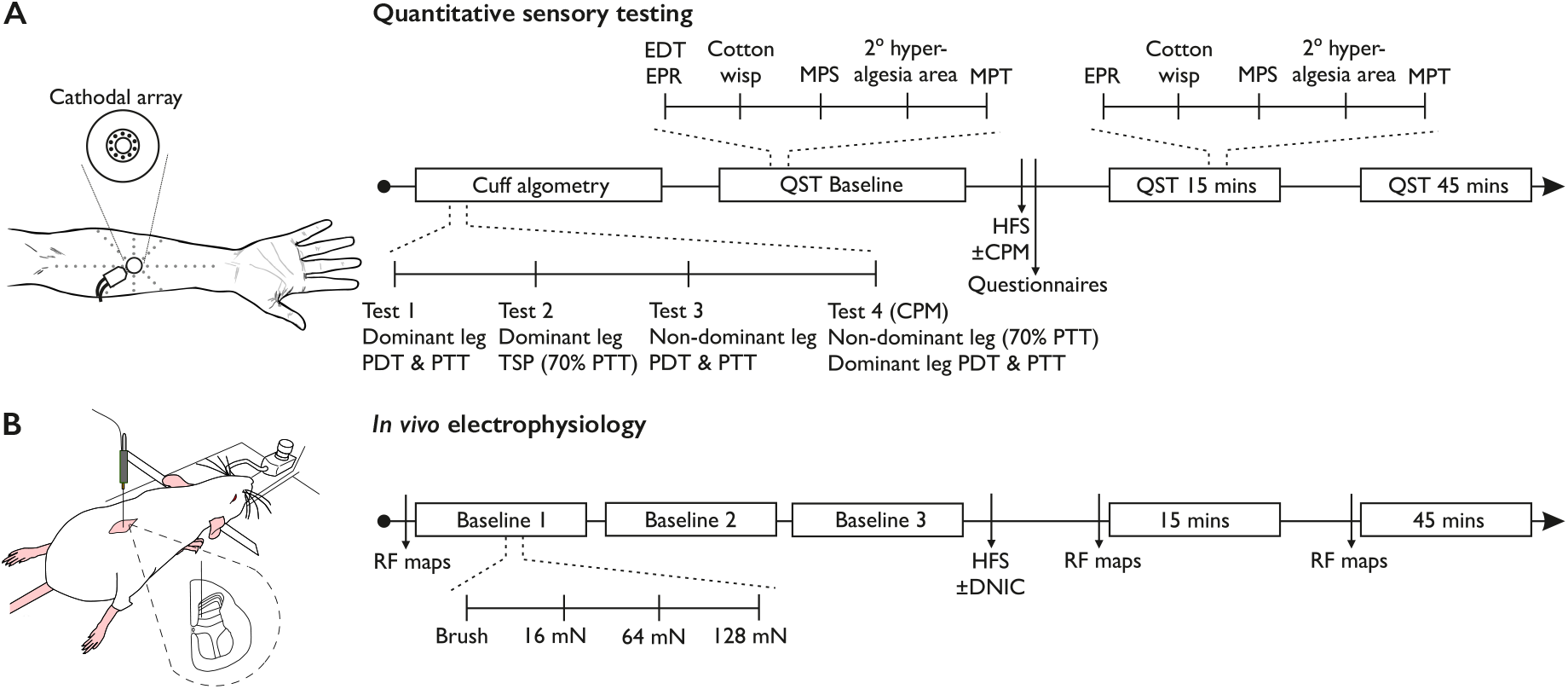
Schematics of experimental sessions. **(A)** Timeline depicting procedure for HFS(control) and HFS(+CPM) quantitative sensory testing sessions in humans. **(B)** Timeline depicting experimental protocol for rat *in vivo* electrophysiology. CPM – conditioned pain modulation, DNIC – diffuse noxious inhibitory controls, EDT – electrical detection threshold, EPR – electrical pain rating, HFS – high frequency stimulation, MPS – mechanical pain sensitivity, MPT – mechanical pain threshold, PDT – pain detection threshold, PTT – pressure tolerance threshold, QST – quantitative sensory testing, RF – receptive field, TSP – temporal summation of pain.

#### 2.2.1 Conditioned pain modulation (CPM) and temporal summation of pain (TSP)

Cuff pressure pain sensitivity and CPM were determined using a computer controlled cuff algometry system (Nocitech CPAR, Aalborg University, Denmark) as previously described (Cummins et al., 2020). In brief, subjects experienced a slow increasing pressure ramp (1 kPa/s up to a maximum of 100 kPa) applied to the calf and were asked to rate pain intensity continuously on a Visual Analogue Scale (VAS) using an electronic VAS device with a sliding bar which digitised to a 0-10 scale. The bar was anchored visually on the controller with ‘min’ and ‘max’, and verbally by the experimenters as ‘no pain at all’ and ‘the worst pain imaginable’. Pain Detection Threshold (PDT) was taken as the pressure (kPa) when a VAS score of 1 out of 10 was reached, and Pain Tolerance Threshold (PTT) was taken when subjects self-terminated the test indicating when they ‘cannot tolerate any more pressure’. For those who reached the maximum of 100 kPa, this was taken as their PTT. Firstly, the pressure ramp was applied to the dominant leg to determine PDT and PTT, followed by assessment of temporal summation of pain (TSP) at a frequency of 1 s on/1 s off (ten stimuli applied at 70% PTT). The pressure ramp was then applied to the non-dominant leg to determine PDT and PTT. To assess CPM, an increasing pressure ramp was applied to the dominant leg and PDT and PTT were determined whilst tonic pressure was applied to the non-dominant leg (70% PTT). The conditioning pressure applied during HFS was taken as 70% of the PTT on the leg contralateral to the subject’s dominant arm, where the electrical stimulation was applied, and was also applied to this leg during the electrical stimulation.

The CPM effect was calculated as the conditioned PDT minus the baseline PDT measurements. This ensured that the presence of an inhibitory CPM effect was indicated by a positive value. The ramps were applied twice at the beginning of each session and the difference was calculated for each session, and then values from each of the two sessions were averaged to calculate the mean CPM effect. Classifying subjects by their CPM response was performed using the Standard Error of Measurement (Cummins et al., 2021; Kennedy et al., 2020) calculated as follows: SEM = Standard deviation of baseline PDT x **√** (1 - intraclass correlation coefficient of baseline PDT). Subjects with a mean CPM response greater than the SEM were classified as CPM responders, those with a response less than the SEM as CPM non-responders. The wind-up ratio (WUR) was calculated as the sum of pain intensity ratings evoked by all ten stimuli divided by the theoretical non-potentiated response (10 x the first stimulus pain intensity rating); values were averaged across the two sessions.

#### 2.2.2 High frequency electrocutaneous stimulation (HFS)

HFS and psychophysical testing was performed as previously described (Klein et al., 2004). Subjects received electrical stimuli via an EPS-P10 electrode (Cathode: 10 pins, 0.25 mm diameter, Anode: 24 × 22 mm^2^; MRC Systems GmbH, Germany), which was attached to the volar part of the subject’s dominant forearm (Fig. 1A). Electrical pulses were delivered transcutaneously via a DS7A High Voltage Constant Current Stimulator (Digitimer Ltd, Welwyn Garden City, UK) whilst trains of stimulation were generated using the DG2A Train Delay Generator (Digitimer Ltd, Welwyn Garden City, UK). Initially following attachment of the electrode, a single 2 ms pulse at 1 mA was administered as a familiarisation to the subject with the electrical stimuli, and to ensure correct placement of the electrode. Subjects then had their electrical detection threshold (EDT) measured using the method of limits. Individual pulses were applied, and stimulus intensity (mA) was gradually decreased until subjects no longer perceived it. The intensity was then increased until the pulses were detected again, and this was repeated five times. The initial value was set at 1 mA to ensure the intensity always began above detection threshold. Non-detection and detection intensities were measured to 0.01 mA resolution and the geometric mean of the ten recorded intensities was taken as the EDT. The intensity of the stimulation was then increased to fifteen times the EDT and was used assess electrical pain ratings (EPR) for the remainder of the session. EDT was measured at the beginning of both sessions, but the intensity used for all electrical stimuli was determined using the EDT from the first session.

Sensitivity to mechanical stimuli was assessed with a shortened version of the DFNS Mechanical Pain Sensitivity (MPS) protocol (Rolke et al., 2006). This utilised only one of the original three non-noxious stimuli (cotton wisp) and four of the pin-prick stimuli (32, 64, 128 and 256 mN; MRC Systems GmbH, Germany), together with single 2 ms electrical pulses applied at 15x EDT. Each stimulus was applied five times within the circular area 1 cm from the edge of the electrode in a counterbalanced pseudo-random order. Subjects were asked rate the painfulness of each stimulus from 0 (no pain at all) to 100 (the worst pain imaginable). Pain ratings for each individual stimulus were averaged both individually and as a whole with the geometric mean of all the pin-prick pain ratings being taken as the subjects’ MPS rating. The full DFNS Mechanical Pain Threshold (MPT) test was also performed. In order to sample all around the electrode, pin-prick stimuli were applied in a clockwise direction around electrode in the same area as the MPS test. The reliability of baseline measures between sessions is shown in Figure S1.

In order to elicit secondary mechanical hyperalgesia, five trains of electrical stimulation (100 Hz, 2 ms pulse width, 1 s duration) were then delivered 10 s apart at 15x EDT. Immediately after the HFS stimulation was applied, subjects completed a measure of both state and trait anxiety (Spielberger et al., 1983) and the SF-36 health questionnaire (Ware and Sherbourne 1992). Sensory testing was then repeated at 15 min and 45 min post-HFS. In order to measure the spread of hyperalgesia, a 10 g von Frey filament (Touch Test, North Coast Medical, Morgan Hill, CA) was applied at 1 cm intervals along eight orthogonal directions around the electrode (Fig. 1A). Subjects were asked to make the same judgement as in the MPT test, and indicate whether the stimulus had a ‘sharp, stinging or pricking sensation’ or a ‘normal touch sensation’. Stimuli were applied from outside to in, starting at a maximum of 8 cm. The spread of hyperalgesia was taken as the number of consecutive ‘sharp’ ratings from the centre (e.g. final three stimulations rated as ‘sharp’ with a ‘blunt’ rating at 4 cm was taken as 3 cm in that direction). The distances from the centre were modelled as an irregular polygon using MATLAB (MathWorks, Natick, MA), the area of which was taken as the area of secondary hyperalgesia.

### 2.3 Animals

Adult male Lister Hooded rats (18 in total; 250-300 g) were used for electrophysiological experiments (Charles River, UK). Animals were group housed (maximum of 5) on a conventional 12 h: 12 h light-dark cycle; food and water were available *ad libitum*. Temperature (20-22 °C) and humidity (55-65 %) of holding rooms were closely regulated. Experimental design/analysis was conducted according to ARRIVE guidelines (appendix 1). All procedures described here were approved by an internal ethics panel and the UK Home Office (licence PABEF3413) under the Animals (Scientific Procedures) Act 1986.

### 2.4 In vivo electrophysiology

Anaesthesia was initially induced with 3.5% v/v isoflurane delivered in 3:2 ratio of nitrous oxide and oxygen. Once areflexic, a tracheotomy was performed and rats were subsequently maintained on 1.5% v/v isoflurane for the remainder of the experiment (approximately 3-4 hours; core body temperature was maintained throughout with the use of a homeothermic blanket). Rats were secured in a stereotaxic frame and a laminectomy was performed to expose the L4-L6 segments of the spinal cord; two spinal clamps were applied to stabilise the spinal column. Extracellular recordings were obtained from deep dorsal horn wide dynamic range lamina V/VI neurones with receptive fields on the glabrous skin of the hind toes using 127 μm diameter 2 MΩ parylene-coated tungsten electrodes (A-M Systems, Sequim, WA). The search stimulus consisted of light tapping of the hind paw as the electrode was manually lowered. Neurones were characterised from depths relating to the deep dorsal horn laminae (HFS 746 ± 100 μm; HFS(+DNIC) 708 ± 66 μm) (Watson et al., 2009), and once a single unit was isolated neurones were classified as wide dynamic range on the basis of sensitivity to dynamic brushing, noxious mechanical (128 mN) and noxious heat stimulation (48 °C) of the receptive field. Data were captured and analysed by a CED Micro1401 interface coupled to a computer with Spike2 v4 software (Cambridge Electronic Design, Cambridge, United Kingdom). The signal was amplified (x7500), bandpass filtered (low/high frequency cut-off 0.5/2 kHz) and digitised at rate of 20 kHz.

DNIC were recruited by applying 40 kPa pressure via a neonatal cuff to the gastrocnemius muscle contralateral to the neuronal recording as previously described (Cummins et al., 2020). HFS was performed in independent experimental groups with and without application of the conditioning stimulus. Electrical stimulation was delivered transcutaneously via needles inserted into the receptive field. Five trains of electrical stimulation (100 Hz, 2 ms pulse width, 1 s duration) were delivered 10 s apart at two times the C-fibre threshold (via a DS3 Isolated Current Stimulator; Digitimer Ltd, Welwyn Garden City, UK). Prior to HFS, receptive field maps were produced in response to 16, 64 and 128 mN pin-prick stimulators (MRC Systems GmbH, Germany). An area was considered part of the receptive field if a response of five action potentials during 1 s stimulation was obtained. A rest period of 20 s between applications was used to avoid sensitisation. Receptive field sizes are expressed as a percentage area of a standardised paw measured using ImageJ (NIH, Bethesda, MD).The receptive field was subsequently stimulated with dynamic brushing (#2 squirrel hair artist’s brush) and 16, 64 and 128 mN pin-prick. Stimuli were applied 50-60 s apart for a duration of 10 s and the evoked response quantified. Baseline data represent mean of three trials performed 5 min apart. Following HFS, receptive field maps and stimulus evoked neuronal responses were determined 15 min and 45 min post-stimulation (Fig. 1B).

### 2.5 Statistics

Statistical analyses were performed using SPSS v28 (IBM, Armonk, NY). The experimental unit for electrophysiological recordings was the individual rat; no animals were excluded from analysis. For psychophysical testing, the experimental unit was the individual subject; forty-five were recruited but eight were excluded from analysis due to withdrawal after the first session (3), incorrect performance with cuff algometry (3), equipment failure (1) and hyposensitivity after HFS (1). Mechanical coding of neurones were compared with a 2-way repeated measures (RM) ANOVA, followed by a Bonferroni *post hoc* test for paired comparisons. Dynamic brush evoked responses were compared with a 1-way RM ANOVA, followed by a Bonferroni *post hoc* test for paired comparisons. Where appropriate, sphericity was tested using Mauchly’s test; the Greenhouse-Geisser correction was applied if violated. Receptive field sizes were compared with a Friedman test followed by Wilcoxon *post hoc* with Bonferroni correction for paired comparisons. For psychophysical measures, data were compared with either a 1-way RM ANOVA or 2-way RM ANOVA followed by a Bonferroni *post hoc* test for paired comparisons. Multivariate linear regression was performed for correlation analysis followed by an ANOVA. Minimum group sizes were determined by *a priori* calculations using the following assumptions (α 0.05, 1-β 0.8, ε 1, effect size range *d*=0.5 to 0.8). Effect sizes for electrophysiology experiments were guided by historical data whereas effect sizes for human psychophysics were based on a pilot study. Each rodent experimental group balanced the need to ensure statistical robustness while adhering to the ‘3 Rs’ (refine, reduce, replace; https://www.nc3rs.org.uk/the-3rs). All data represent mean ± 95% confidence interval (CI). * *P*<0.05, \*\**P*<0.01, \*\*\**P*<0.001.

## 3. Results

### 3.1 High frequency electrocutaneous stimulation induces secondary but not primary hyperalgesia in humans

During high frequency electrocutaneous stimulation, a temporal summation effect was observed with pain intensity ratings higher during the fifth stimulus train compared to the first (1 way RM ANOVA, *F*_1.38,49.73_=53.18, *P*=5.23×10^−11^) (Fig. 2A). As a measure of primary hyperalgesia, electrical pain ratings (EPR) were compared prior to and post-HFS. Primary hyperalgesia was reported by 9/37 subjects and overall no time dependent change in pain intensity ratings was observed (1 way RM ANOVA, *F*_1.55,55.62_=1.356, *P*=0.267) (Fig. 2B). Secondary brush allodynia was infrequent and only reported by 3/37 subjects (1 way RM ANOVA, *F*_1.54,55.29_=2.423, *P*=0.111) (Fig. 2C). Secondary pin-prick hyperalgesia was more a prominent feature of sensitisation as evidenced by an increase in the area of secondary hyperalgesia in 28/37 subjects (1 way RM ANOVA, *F*_2,72_=21.787, *P*=3.99×10^−8^) (Fig. 2D), an increase in mechanical pain sensitivity in 35/37 subjects (1 way RM ANOVA, *F*_1.60,57.42_=16.156, *P*=0.00001) (Fig. 2E), and a decrease in mechanical pain threshold in 33/37 subjects (1 way RM ANOVA, *F*_1.18,42.42_=19.226, *P*=0.00003) (Fig. 2F). Pain intensity ratings to increasing pin-prick forces exhibited a linear stimulus-response relationship and were increased at both time-points post-HFS (2 way RM ANOVA, time x stimulus intensity interaction *F*_3.43,123.4_=5.196, *P*=0.00005) (Fig. 2F).

**Figure 2.**
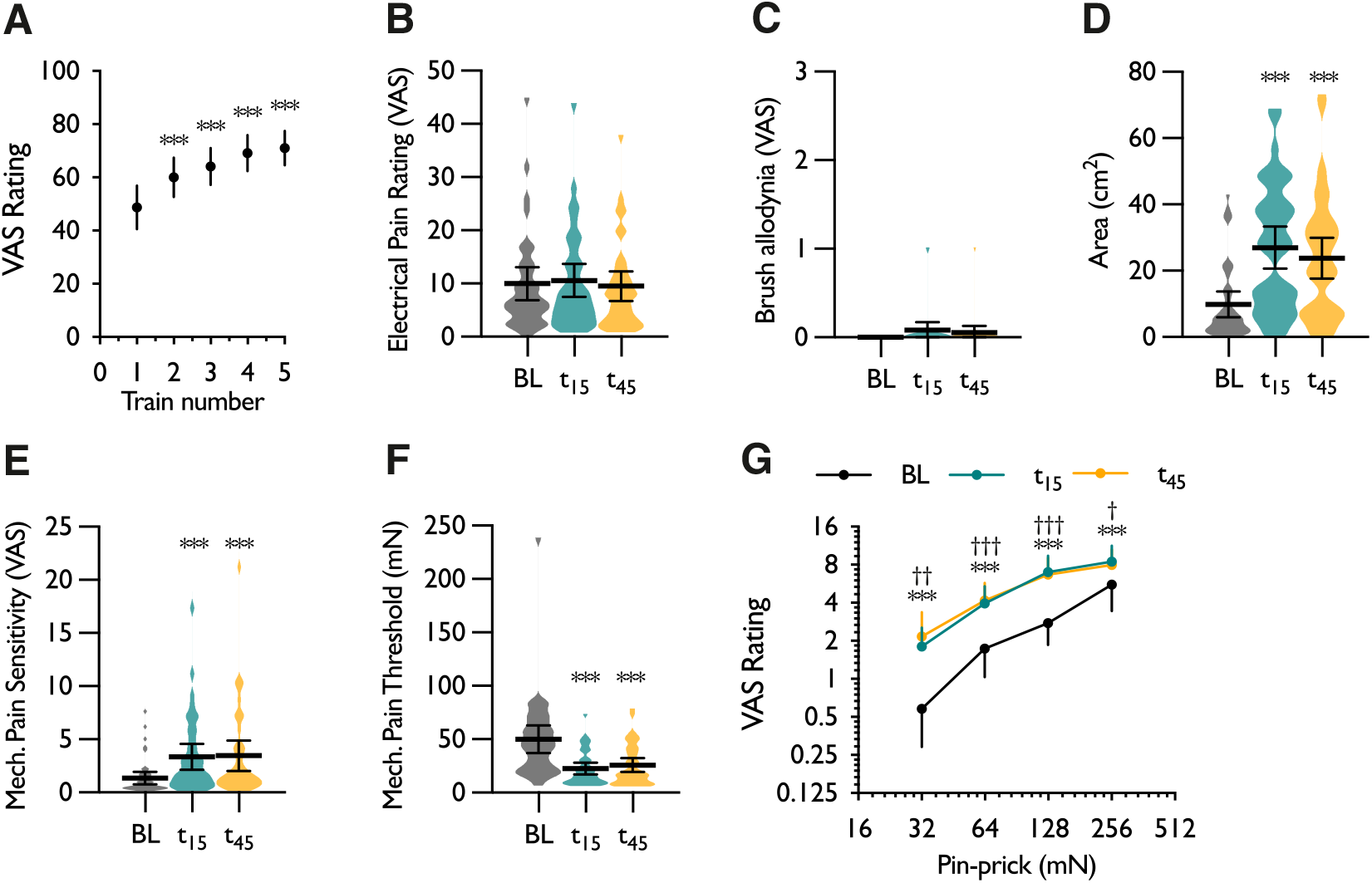
Primary and secondary measures of sensitisation following high frequency electrocutaneous stimulation. **(A)** Pain intensity ratings to repeated trains of electrical stimulation. **(B)** Pain intensity ratings to a single pulse electrical stimulus (primary hyperalgesia). **(C)** Secondary brush allodynia. **(D)** Area of secondary hyperalgesia in response to a 10 g von Frey filament. **(E)** Secondary mechanical pain sensitivity. **(F)** Secondary mechanical pain threshold. **(G)** Pain intensity ratings to individual pin-prick forces. Data represent mean ± 95% CI; *n*=37. \**P*<0.05, \*\**P*<0.01, \*\*\**P*<0.001; For panels A-E, * denotes difference from baseline. For panel G, * denotes difference between baseline and 15 min time-point (t_15_), † denotes difference between baseline and 45 min time-point (t_45_). BL – baseline, VAS – visual analogue scale.

Cuff pressure algometry was performed at baseline to assess temporal summation of pain and a progressive increase in the pain intensity rating from the first to tenth stimulus was observed (1 way RM ANOVA, *F*_2.31,83.01_=34.3, *P*= 1.48×10^−12^) (Fig. 3A). The wind-up ratio was calculated for both cuff algometry and electrical stimulation and no correlation between the two was found (Fig. 3B). In addition we examined whether wind-up ratios or CPM efficiency correlated with the degree of secondary hyperalgesia. The pressure/electrical wind-up ratios and the CPM effect did not correlate with a change in the area of secondary hyperalgesia (adj. R^2^=-0.099; ANOVA *F*_3,28_=0.067, *P*=0.977) (Fig. 3C-E), or with the change in mechanical pain sensitivity (adj. R^2^=0.037; ANOVA *F*_3,28_=1.402, *P*=0.263) (Fig. 3F-H), or with the change in mechanical pain threshold (adj. R^2^=-0.08; ANOVA *F*_3,28_=0.235, *P*=0.872) (Fig. 3I-K). The development of secondary hyperalgesia was also not dependent on HFS intensity (mean 1.70 ± 0.35 mA) (Table S1), or related to general health and state trait anxiety scores (Table S2).

**Figure 3.**
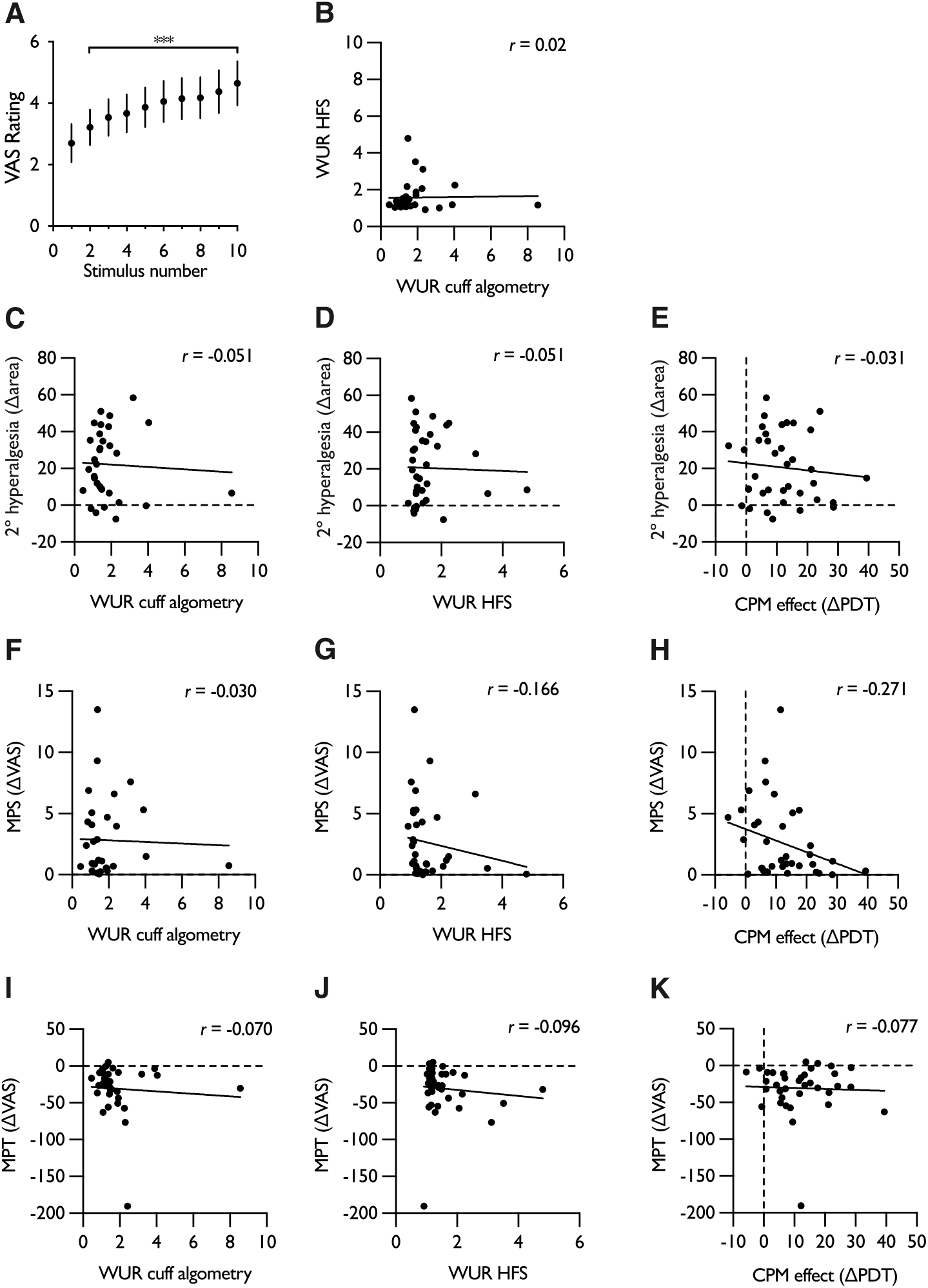
Correlation of wind-up ratios and CPM efficiency with measures of secondary hyperalgesia. **(A)** Temporal summation of pain in response to repetitive cuff pressure stimulation. **(B)** Correlation of electrical and cuff algometry wind-up ratios. **(C)** Correlation of cuff algometry wind-up ratio with area of secondary hyperalgesia. **(D)** Correlation of HFS wind-up ratio with area of secondary hyperalgesia. **(E)** Correlation of CPM effect with area of secondary hyperalgesia. **(F)** Correlation of cuff algometry wind-up ratio with mechanical pain sensitivity. **(G)** Correlation of HFS wind-up ratio with mechanical pain sensitivity. **(H)** Correlation of CPM effect with mechanical pain sensitivity. **(I)** Correlation of cuff algometry wind-up ratio with mechanical pain threshold. **(J)** Correlation of HFS wind-up ratio with mechanical pain threshold. **(K)** Correlation of CPM effect with mechanical pain threshold. For all dependent measures, peak change from baseline is used for comparison. For panel A, data represent mean ± 95% CI; *n*=32, \*\*\**P*<0.001; * denotes difference from first stimulus. CPM – conditioned pain modulation, HFS – high frequency stimulation, MPS – mechanical pain sensitivity, VAS – visual analogue scale, WUR – wind-up ratio.

### 3.2 Applying a conditioned pain modulation paradigm concurrent to high frequency stimulation does not alter the development of secondary hyperalgesia in humans

CPM was assessed with cuff pressure algometry to stratify the initial cohort (Fig. 4A). Based on a standard error of measurement of 8.64, subjects reporting an increase in pain detection threshold above this were classified as CPM responders (21/37). In this sub-group, concurrently applying a heterotopic noxious conditioning stimulus during high frequency stimulation did not affect reported pain intensity ratings to the five electrical trains (2 way RM ANOVA, stimulus x CPM interaction *F*_2.65,53.07_=0.241, *P*=0.845) (Fig. 4B). In the HFS control session, a time-dependent decrease in the primary pain intensity rating was observed at the 45 min time-point (1 way RM ANOVA, *F*_2,40_=4.74, *P*=0.018), however there was no change in the HFS(+CPM) session (1 way RM ANOVA, *F*_1.28,25.75_=0.332, *P*=0.625) (Fig. 4C). Comparing the interactive effect between experimental sessions revealed a difference in baseline electrical pain ratings (2 way RM ANOVA, time x CPM interaction *F*_2.,40_=4.393, *P*=0.019) (Fig. 4C). A time-dependent increase in the area of secondary hyperalgesia was observed in both the HFS control (1 way RM ANOVA, *F*_1.57,31.42_=14.193, *P*=0.0001) and HFS(+CPM) sessions (1 way RM ANOVA, *F*_2,40_=10.97, *P*=0.00016), however CPM had no effect on the spread of secondary hyperalgesia (2 way RM ANOVA, time x CPM interaction *F*_2.,40_=0.527, *P*=0.594) (Fig. 4D). A time-dependent increase in mechanical pain sensitivity was observed in both the HFS control (1 way RM ANOVA, *F*_1.49,29.87_=6.539, *P*=0.0082) and HFS(+CPM) sessions (1 way RM ANOVA, *F*_1.4,28.00_=6.312, *P*=0.011), however CPM had no effect on mechanical pain sensitivity (2 way RM ANOVA, time x CPM interaction *F*_2.,40_=0.581, *P*=0.564) (Fig. 4E). A time-dependent decrease in mechanical pain threshold was observed in both the HFS control (1 way RM ANOVA, *F*_1.13,22.57_=8.828, *P*=0.0055) and HFS(+CPM) sessions (1 way RM ANOVA, *F*_1.06,21.22_=6.283, *P*=0.019), however CPM had no effect on mechanical pain thresholds (2 way RM ANOVA, time x CPM interaction *F*_1.15.,22.99_=0.09, *P*=0.914) (Fig. 4F). When comparing the peak change in pain intensity ratings to individual pin-prick forces, we found no evidence for an effect of CPM during HFS (3 way ANOVA, time x CPM x stimulus intensity interaction *F*_3,120_=1.112, *P*=0.347) (Fig. 4G).

**Figure 4.**
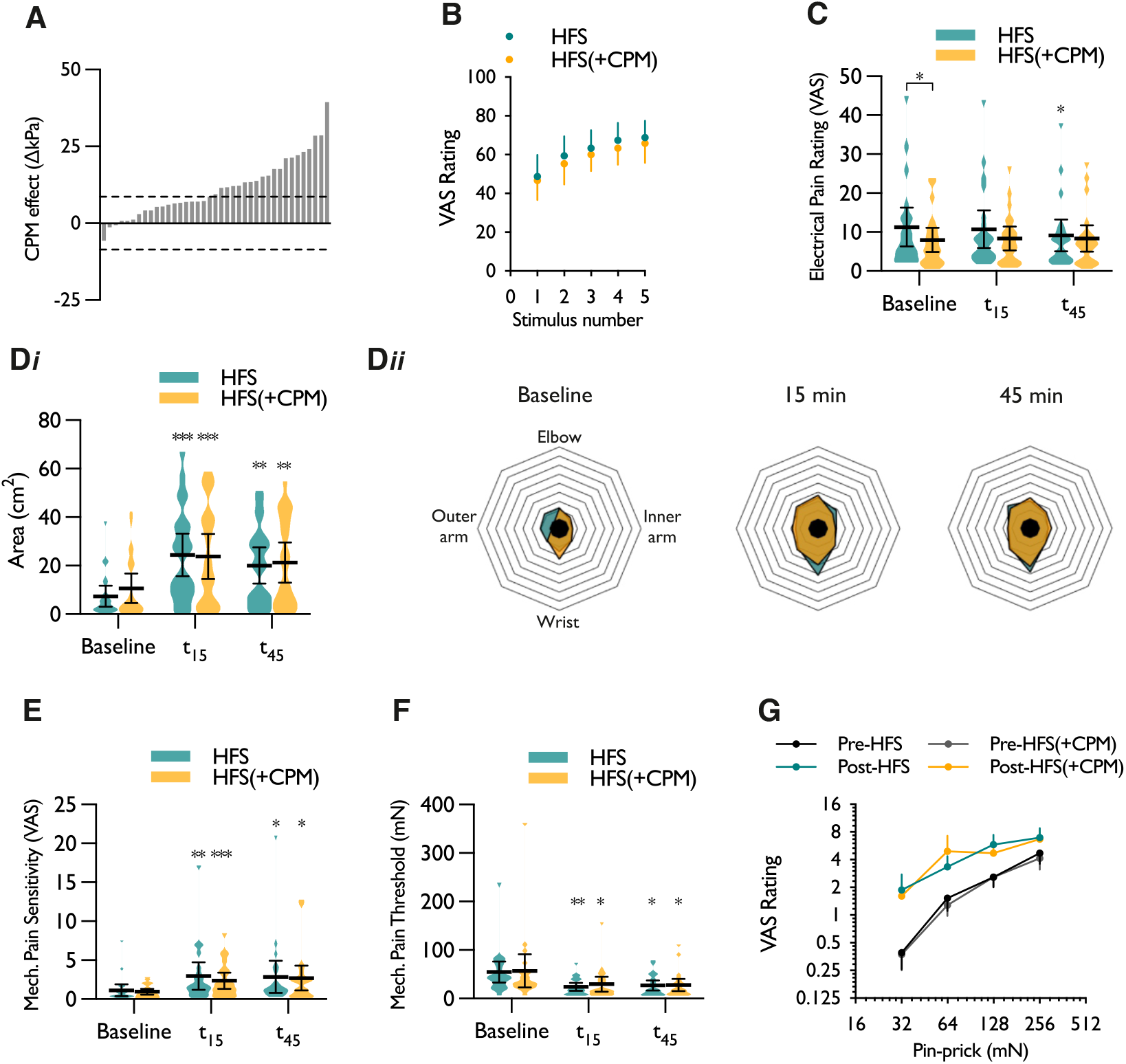
Effect of conditioned pain modulation on the development of electrically induced secondary hyperalgesia. **(A)** Stratification of subjects based on CPM effect determined by a change in pain detection threshold. Bars represent individual responses, dashed lines represent standard error of measurement. **(B)** Pain intensity ratings to repeated trains of electrical stimulation with and without CPM applied. **(C)** Pain intensity ratings to a single pulse electrical stimulus (primary hyperalgesia). **(D*i*)** Area of secondary hyperalgesia in response to a 10 g von Frey filament, and **(D*ii*)** polygon plots of the mean spread of all subjects across experimental sessions. **(E)** Secondary mechanical pain sensitivity. **(F)** Secondary mechanical pain threshold. **(G)** Peak change of pain intensity rating to individual pin-prick forces. Data represent mean ± 95% CI; *n*=21. \**P*<0.05, \*\**P*<0.01, \*\*\**P*<0.001; Unless otherwise indicated, * denotes difference between time-point and respective baseline.

### 3.3 Activation of diffuse noxious inhibitory controls does not supress the development of high frequency stimulation-induced neuronal sensitisation in rats

Electrical stimulation was delivered transcutaneously in rats using identical parameters to the human study (Fig. 5A). The stimulation intensity was comparable between experimental groups (HFS: 1.76 ± 0.73 mA, HFS(+DNIC): 1.18 ± 0.28 mA; unpaired T test with Welch’s correction, *P*=0.176). HFS produced a transient primary brush hypersensitivity in the absence of a conditioning stimulus (1 way RM ANOVA, *F*_2,16_=6.485, *P*=0.0087) (Fig. 5B), whereas brush hypersensitivity was less pronounced after HFS when performed concurrently with tonic pressure applied to the contralateral leg (1 way RM ANOVA, *F*_1.093,8.745_=2.926, *P*=0.121) (Fig. 5C). Compared to brush hypersensitivity, HFS produced a longer lasting primary pin-prick hypersensitivity to range of non-noxious and noxious intensities of stimulation (2 way RM ANOVA, time x stimulus intensity interaction *F*_4,32_=2.811, *P*=0.042) (Fig. 5D). Pin-prick hypersensitivity was still robustly induced following HFS performed while DNIC were active (2 way RM ANOVA, time x stimulus intensity interaction *F*_4,32_=3.805, *P*=0.012) (Fig. 5E), however when comparing the peak change from baseline no differences in pin-prick hypersensitivity were observed between the control and DNIC experiments (3 way ANOVA, HFS x DNIC x stimulus intensity interaction *F*_2,32_=0.390, *P*=0.684) (Fig. 5F). An expansion of receptive field size was observed following HFS with increases in response to 16 mN (Friedman’s test, *P*=0.006), 64 mN (Friedman’s test, *P*=0.00086) and 128 mN (Friedman’s test, *P*=0.00049) pin-prick stimuli (Fig. 5G). In the HFS(+DNIC) experiment, the expansion of receptive fields in response to 16 mN stimulation was less pronounced and weak evidence was found for increased responsivity (Friedman’s test, *P*=0.044; paired comparisons *P*>0.05), however increased responsivity to 64 mN (Friedman’s test, *P*=0.004) and 128 mN (Friedman’s test, *P*=0.0037) stimuli was still observed (Fig. 5H) of similar magnitude to the control experiment (mean fold increase in RF size: HFS_15min_ and HFS(+DNIC)_15min_ – 2.04 and 1.88 (16 mN), 2.62 and 1.66 (64 mN), 2.15 and 1.93 (128 mN)).

**Figure 5.**
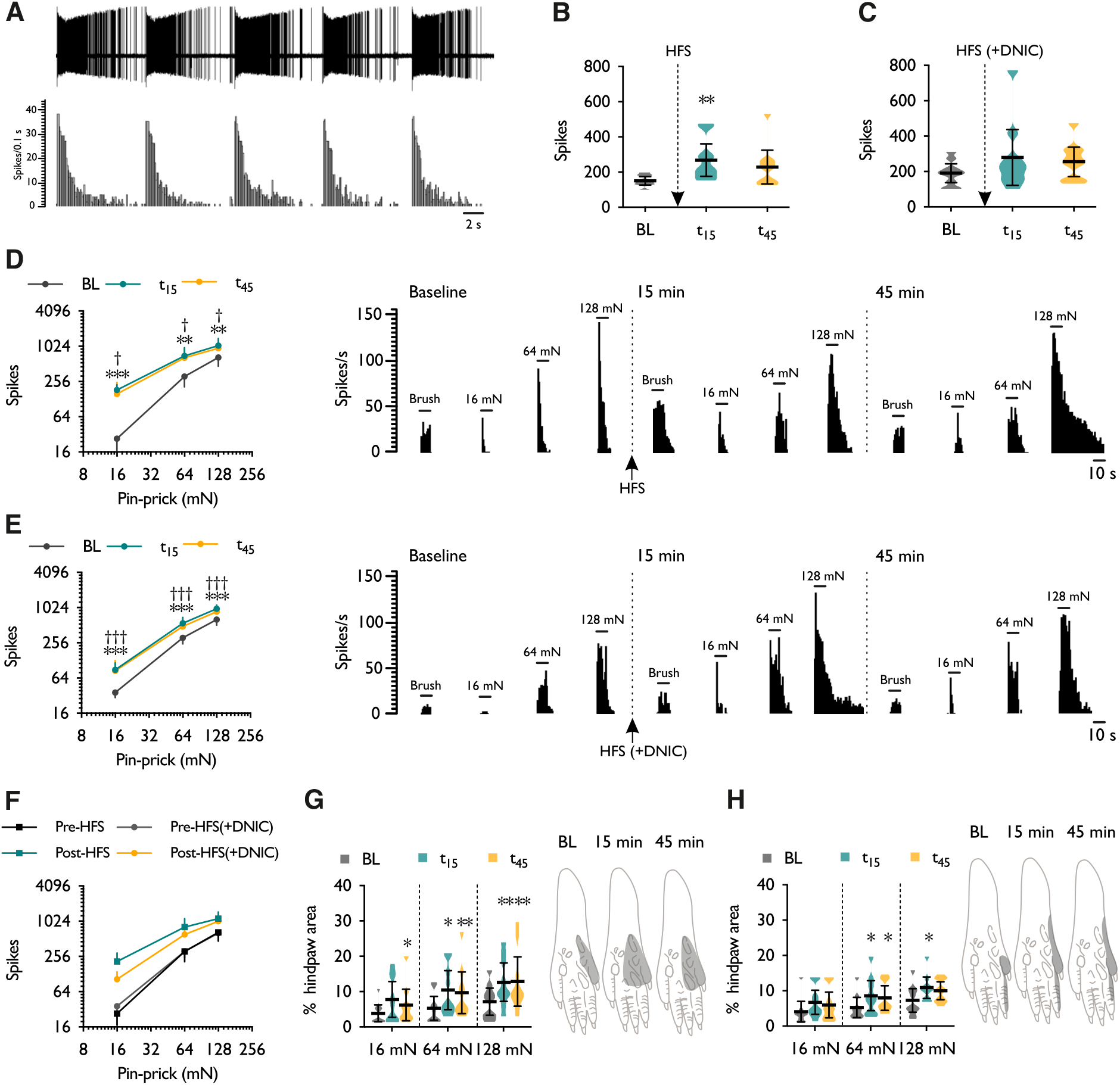
Effect of activating diffuse noxious inhibitory controls on the development of electrically induced spinal neuronal hyperexcitability. **(A)** Representative neurogram and histogram trace of single unit response to high frequency electrocutaneous stimulation. Dynamic brush evoked neuronal responses prior to- and post-electrical stimulation in the absence and **(C)** presence of a conditioning stimulus. Pin-prick evoked neuronal responses prior to- and post-electrical stimulation in the **(D)** absence and **(E)** presence of a conditioning stimulus. Receptive field sizes to pin-prick stimuli prior to- and post-electrical stimulation in the **(F)** absence and **(G)** presence of a conditioning stimulus. Schematics of hind paws represent typical single unit receptive field to 128 mN stimulation. Data represent mean ± 95% CI; *n*_HFS_=9, *n*_HFS(+DNIC)_=9. \**P*<0.05, \*\**P*<0.01, \*\*\**P*<0.001; * denotes difference between baseline and 15 min time-point (t_15_), † denotes difference between baseline and 45 min time-point (t_45_). DNIC – diffuse noxious inhibitory controls, HFS – high frequency stimulation.

## 4. Summary

- The parallel human and rat study design provides insight into the input-output relationship between spinal neuronal activity and perceptual responses. Human psychophysics and rat spinal neurones display similar stimulus-response relationships to pin-prick stimulation prior to and post-electrical conditioning. This alignment supports the translational value of the pre-clinical model to assess neural substrates of hyperalgesia.
- We did not find a correlation between wind-up ratios during HFS and during cuff algometry, which could be attributed to the nature of the stimuli. Electrical stimulation was delivered cutaneously at suprathreshold levels of stimulation whereas pressure stimulation during cuff algometry assessed temporal summation of deep inputs at threshold levels potentially recruiting different amplification mechanisms. We did not find that baseline temporal summation of pain or CPM efficiency explained the variability of secondary hyperalgesia elicited.
- CPM had no effect on the development of secondary hyperalgesia or on electrically evoked pain ratings during HFS. An effective CPM response was confirmed at baseline using cuff algometry and the lack of interaction with electrical stimulation could be explained by the intensity of the conditioning stimulus suggesting at this level of conditioning excitatory drive exceeds inhibitory controls. Although DNIC and CPM are not equivalent measures, we found a concordance between human and rat data as DNIC had minimal effect on HFS-induced neuronal sensitisation when applying comparable paradigms.

## Abbreviations

CPM: conditioned pain modulation,
DNIC: diffuse noxious inhibitory controls,
EDT: electrical detection threshold,
EPR: electrical pain rating,
HFS: high frequency stimulation,
LTP: long term potentiation,
MPS: mechanical pain sensitivity,
MPT: mechanical pain threshold,
PDT: pain detection threshold,
PTT: pressure tolerance threshold,
QST: quantitative sensory testing,
RF: receptive field,
TSP: temporal summation of pain,
VAS: visual analogue scale,
WUR: wind-up ratio.

## Acknowledgements

The authors would like to recognise the contribution of Tatum Cummins to the initial piloting of the high frequency stimulation paradigm, and additionally thank Michael Mansfield for assisting.

## Author contributions

RP, JT, AHD, SBM, KB - design of study; AHD, SBM, KB - provided analytical tools and equipment; RP, JT - performed experiments; RP, JT - analysed data; RP, JT, AHD, KB - interpreted results of experiments; RP, JT - drafted manuscript; RP, JT, AHD, KB - edited and revised manuscript.

## Conflict of interest

The authors have no conflicts of interest to declare.

**Supplementary Figure S1.**
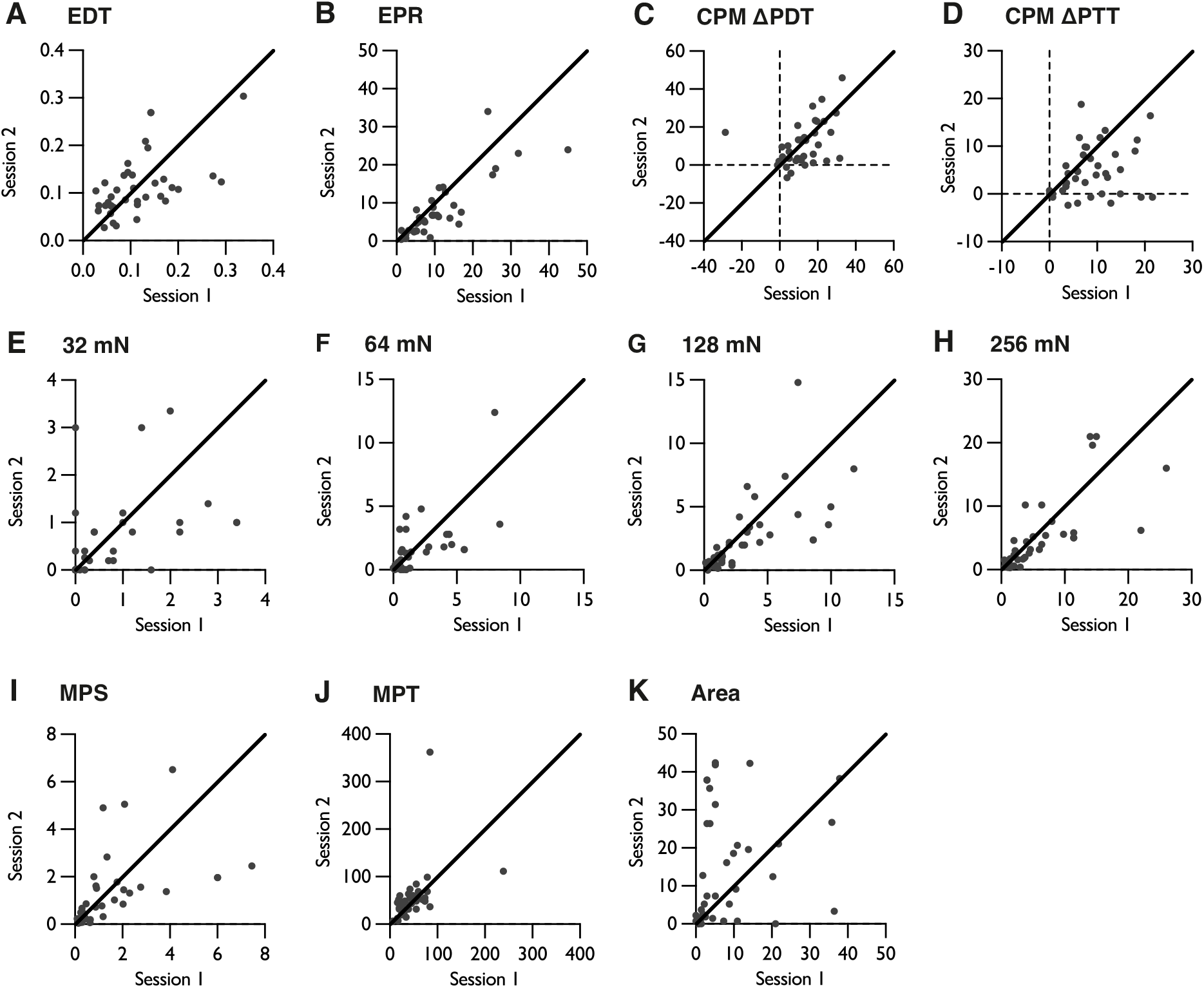
Correlation of baseline measures in session 1 and session 2. **(A)** Electrical detection threshold (EDT) *r*=0.548**. **(B)** Electrical pain rating (EPR) to a single electrical pulse *r*=0.835**. **(C)** Conditioned pain modulation (CPM) effect on pain detection threshold (PDT) *r*=0.423**. **(D)** CPM effect on pain tolerance threshold (PTT) *r*=0.272. **(E)** Pain intensity rating to 32 mN pin-prick *r*=0.835**. **(F)** Pain intensity rating to 64 mN pin-prick *r*=0.680*. **(G)** Pain intensity rating to 128 mN pin-prick *r*=0.685**. **(H)** Pain intensity rating to 256 mN pin-prick *r*=0.745**. **(I)** Mechanical pain sensitivity (MPS) *r*=0.509**. **(J)** Mechanical pain threshold (MPT) *r*=0.414**. **(K)** Area of sensitivity to 10 g von Frey stimulation *r*=0.210. The black line indicates *y*=*x* and represents perfect equivalence between the sessions; *n*=37, \*\**P*<0.01.

**Supplementary Table S1.** Correlations between the intensity of the HFS trains (mA) and the mean pain intensity ratings for five electrical HFS trains (VAS) with each of the dependant measures at each timepoint (calculated as difference from baseline) in the HFS(control) session; *n*=37, \**P*<0.05, \*\**P*<0.01. HFS – high frequency stimulation, MPS – mechanical pain sensitivity, MPT – mechanical pain threshold.

|  | *r* = | HFS Intensity (mA) | Mean HFS(control) VAS |
| --- | --- | --- | --- |
| Area 15 min |  | 0.240 | 0.294 |
| Area 45 min |  | 0.386* | 0.302 |
| MPS 15 min |  | -0.0941 | 0.524** |
| MPS 45 min |  | -0.0244 | 0.461** |
| MPT 15 min |  | -0.218 | -0.116 |
| MPT 45 min |  | -0.191 | -0.0886 |

**Supplementary Table S2.**
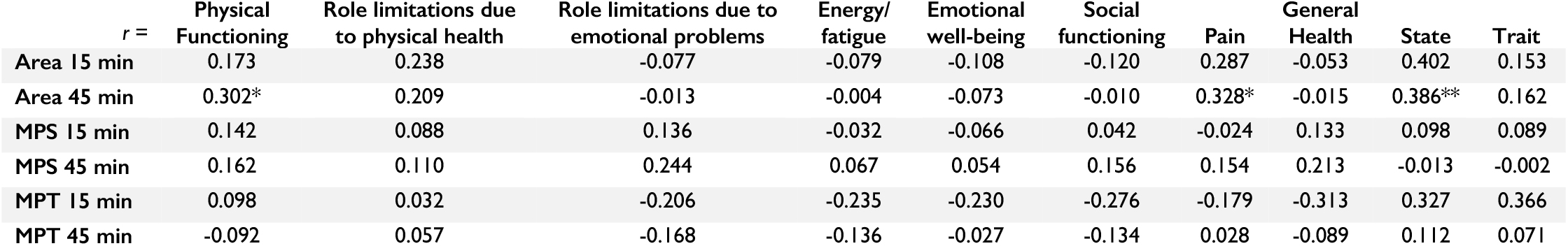
Correlations between scores for general health and state trait anxiety with each of the dependant measures at each timepoint (calculated as difference from baseline) in the HFS(control) session; *n*=37, \**P*<0.05, \*\**P*<0.01. HFS – high frequency stimulation, MPS – mechanical pain sensitivity, MPT – mechanical pain threshold.

| *r* = | Physical<br>Functioning | Role limitations due<br>to physical health | Role limitations due to<br>emotional problems | Energy/<br>fatigue | Emotional<br>well-being | Social<br>functioning | Pain | General<br>Health | State | Trait |
| --- | --- | --- | --- | --- | --- | --- | --- | --- | --- | --- |
| Area 15 min | 0.173 | 0.238 | -0.077 | -0.079 | -0.108 | -0.120 | 0.287 | -0.053 | 0.402 | 0.153 |
| Area 45 min | 0.302* | 0.209 | -0.013 | -0.004 | -0.073 | -0.010 | 0.328* | -0.015 | 0.386** | 0.162 |
| MPS 15 min | 0.142 | 0.088 | 0.136 | -0.032 | -0.066 | 0.042 | -0.024 | 0.133 | 0.098 | 0.089 |
| MPS 45 min | 0.162 | 0.110 | 0.244 | 0.067 | 0.054 | 0.156 | 0.154 | 0.213 | -0.013 | -0.002 |
| MPT 15 min | 0.098 | 0.032 | -0.206 | -0.235 | -0.230 | -0.276 | -0.179 | -0.313 | 0.327 | 0.366 |
| MPT 45 min | -0.092 | 0.057 | -0.168 | -0.136 | -0.027 | -0.134 | 0.028 | -0.089 | 0.112 | 0.071 |

## Appendix 1

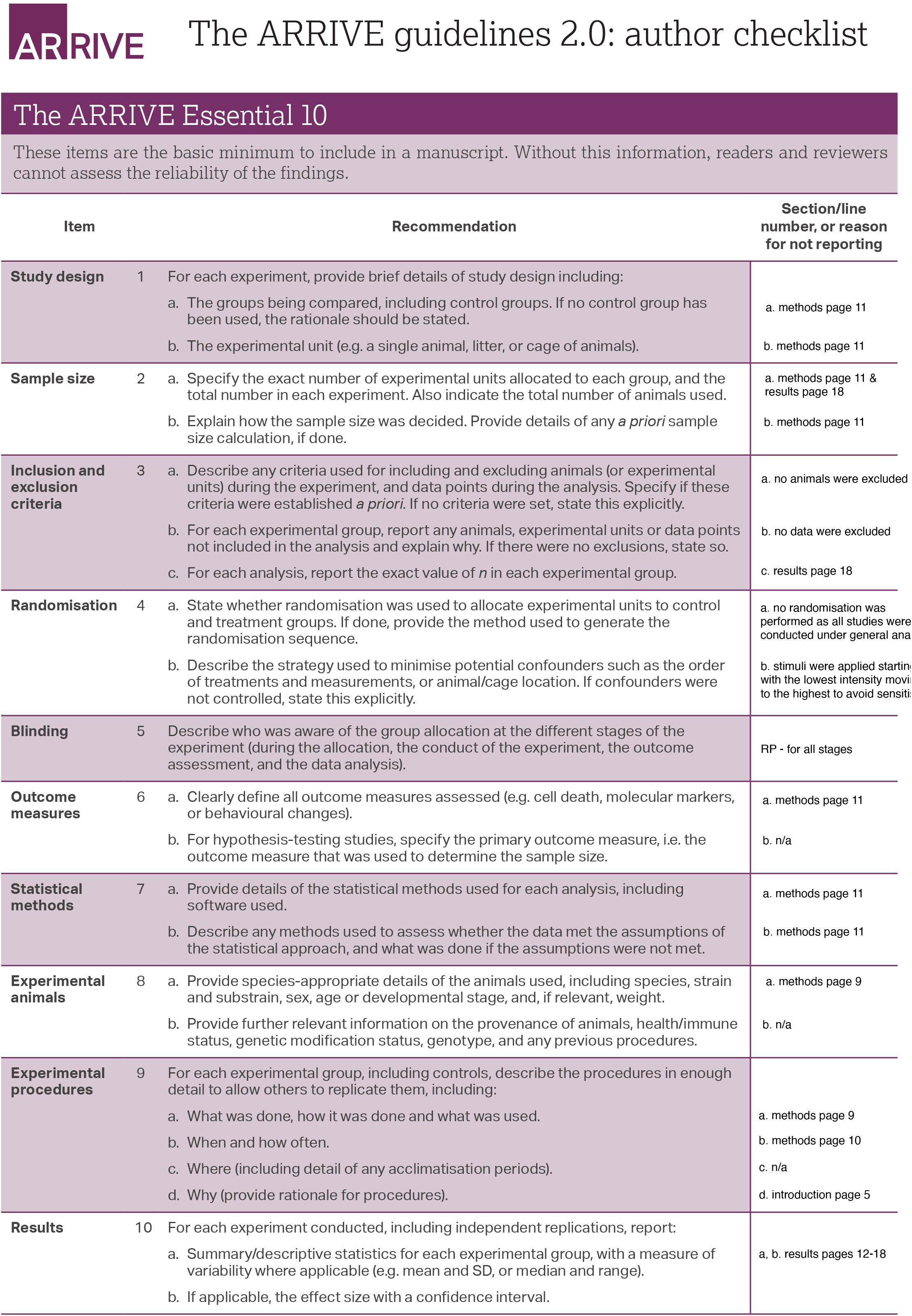

## References

Bannister K, Patel R, Goncalves L, Townson L, Dickenson AH. Diffuse noxious inhibitory controls and nerve injury: restoring an imbalance between descending monoamine inhibitions and facilitations. Pain 2015;156: 1803–1811.

Coghill RC, Mayer DJ, Price DD. Wide dynamic range but not nociceptive-specific neurons encode multidimensional features of prolonged repetitive heat pain. Journal of neurophysiology 1993;69: 703–716.

Cummins TM, Kucharczyk MM, Graven-Nielsen T, Bannister K. Activation of the descending pain modulatory system using cuff pressure algometry: Back translation from man to rat. European journal of pain (London, England) 2020;24: 1330–1338.

Cummins TM, McMahon SB, Bannister K. The impact of paradigm and stringent analysis parameters on measuring a net conditioned pain modulation effect: A test, retest, control study. European journal of pain (London, England) 2021;25: 415–429.

Gjerstad J, Tjølsen A, Hole K. Induction of long-term potentiation of single wide dynamic range neurones in the dorsal horn is inhibited by descending pathways. Pain 2001;91: 263–268.

Henrich F, Magerl W, Klein T, Greffrath W, Treede RD. Capsaicin-sensitive C- and A-fibre nociceptors control long-term potentiation-like pain amplification in humans. Brain 2015;138: 2505–2520.

Ikeda H, Stark J, Fischer H, Wagner M, Drdla R, Jäger T, Sandkühler J. Synaptic amplifier of inflammatory pain in the spinal dorsal horn. Science 2006;312: 1659–1662.

Kennedy DL, Kemp HI, Wu C, Ridout DA, Rice ASC. Determining Real Change in Conditioned Pain Modulation: A Repeated Measures Study in Healthy Volunteers. The journal of pain : official journal of the American Pain Society 2020;21: 708–721.

Klein T, Magerl W, Hopf HC, Sandkühler J, Treede RD. Perceptual correlates of nociceptive long-term potentiation and long-term depression in humans. The Journal of neuroscience : the official journal of the Society for Neuroscience 2004;24: 964–971.

Kucharczyk MW, Di Domenico F, Bannister K. The origin of diffuse noxious inhibitory controls. bioRxiv 2022: 2022.2003.2029.486214.

Le Bars D, Dickenson AH, Besson JM. Diffuse noxious inhibitory controls (DNIC). I. Effects on dorsal horn convergent neurones in the rat. Pain 1979a;6: 283–304.

Le Bars D, Dickenson AH, Besson JM. Diffuse noxious inhibitory controls (DNIC). II. Lack of effect on non-convergent neurones, supraspinal involvement and theoretical implications. Pain 1979b;6: 305–327.

Liu XG and Sandkühler J. Long-term potentiation of C-fiber-evoked potentials in the rat spinal dorsal horn is prevented by spinal N-methyl-D-aspartic acid receptor blockage. Neurosci Lett 1995;191: 43–46.

Maixner W, Dubner R, Bushnell MC, Kenshalo DR, Oliveras JL. Wide-Dynamic-Range Dorsal Horn Neurons Participate in the Encoding Process by Which Monkeys Perceive the Intensity of Noxious Heat Stimuli. Brain Research 1986;374: 385–388.

Mendell LM and Wall PD. Responses of Single Dorsal Cord Cells to Peripheral Cutaneous Unmyelinated Fibres. Nature 1965;206: 97–99.

O’Neill J, Sikandar S, McMahon SB, Dickenson AH. Human psychophysics and rodent spinal neurones exhibit peripheral and central mechanisms of inflammatory pain in the UVB and UVB heat rekindling models. J Physiol 2015;593: 4029–4042.

Quesada C, Kostenko A, Ho I, Leone C, Nochi Z, Stouffs A, Wittayer M, Caspani O, Brix Finnerup N, Mouraux A, Pickering G, Tracey I, Truini A, Treede R-D, Garcia-Larrea L. Human surrogate models of central sensitization: A critical review and practical guide. European Journal of Pain 2021;25: 1389–1428.

Randić M, Jiang MC, Cerne R. Long-term potentiation and long-term depression of primary afferent neurotransmission in the rat spinal cord. The Journal of neuroscience : the official journal of the Society for Neuroscience 1993;13: 5228–5241.

Rolke R, Magerl W, Campbell KA, Schalber C, Caspari S, Birklein F, Treede R-D. Quantitative sensory testing: a comprehensive protocol for clinical trials. European Journal of Pain 2006;10: 77–77.

Sikandar S, Ronga I, Iannetti GD, Dickenson AH. Neural coding of nociceptive stimuli-from rat spinal neurones to human perception. Pain 2013;154: 1263–1273.

Spielberger C, Gorsuch R, Lushene R, Vagg PR, Jacobs G. Manual for the State-Trait Anxiety Inventory (Form Y1 – Y2). 1983.

Ware JE, Jr. and Sherbourne CD. The MOS 36-item short-form health survey (SF-36). I. Conceptual framework and item selection. Med Care 1992;30: 473–483.

Watson C, Paxinos G, Kayalioglu G, Heise C. Chapter 15 - Atlas of the Rat Spinal Cord. In: The Spinal Cord. San Diego: Academic Press; 2009; 238–306.

WMA. World Medical Association Declaration of Helsinki: ethical principles for medical research involving human subjects. Jama 2013;310: 2191–2194.

