## Supplementary figures and images for "A back-translational study of descending interactions with central mechanisms of hyperalgesia induced by high frequency stimulation in rat and human"

### Supplementary figure 1

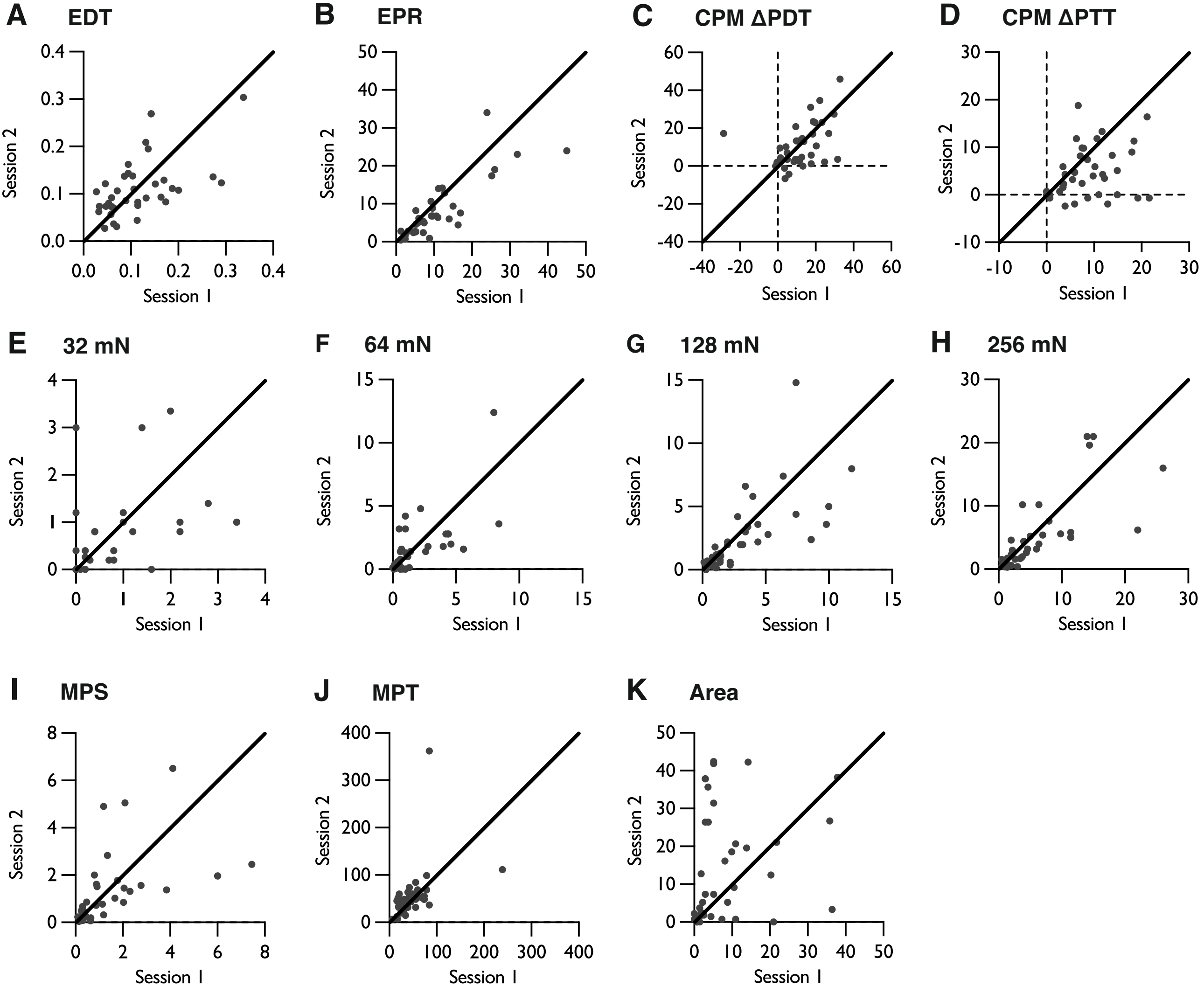
